# Can immunosuppressed mice control oral infection by the opportunistic pathogen *Encephalitozoon intestinalis*?

**DOI:** 10.1101/611988

**Authors:** Maria Lucia da Costa Moura, Anuska Marcelino Alvares-Saraiva, Elizabeth Cristina Pérez, José Guilherme Xavier, Diva Denelle Spadacci-Morena, Carla Renata Serantoni Moysés, Paulo Ricardo Dell’Armelina Rocha, Maria Anete Lallo

## Abstract

Intestinal mucosa (IM), or the outer surface of the intestine, serves at the primary site for the interaction of various pathogens that cause infection via the oral route. Thus, IM is crucial for developing an efficient adaptive immune response against pathogenic micro-organisms, thereby preventing their colonization and subsequent infection. In the present study, we investigated the immune response to *Encephalitozoon intestinalis*-caused infection in the IM and gut-associated lymphoid tissue (GALT) in C57BL/6 female mice. To mimic an immunosuppressive condition, the mice were treated with cyclophosphamide (Cy). Histopathology revealed lymphoplasmacytic enteritis at 7 and 14 days-post-infection (dpi) in all infected groups; however, inflammation diminished at 21 and 28 dpi. Cy treatment also led to a higher number of *E. intestinalis* spores and lesions, which reduced at 28 dpi. In addition, flow cytometry analysis demonstrated CD4^+^ and CD8^+^ T cells to be predominant immune cells, with a significant increase in both Th1 and Th2 cytokines at 7 and 14 dpi, as demonstrated by histopathology. In conclusion, Cy treatment reduced GALT (Peyer’s plaques and mesenteric lymph nodes) and peritoneum populations but increased the T-cell population in the intestinal mucosa and the production of pro-inflammatory cytokines, which were able to eliminate this opportunistic fungus and reduced the infection.

## Introduction

Microsporidia are spore-forming intracellular pathogens, responsible for causing opportunistic infections in immunocompromised people, such as those with HIV infection and AIDS, cancer patients or individuals with autoimmune diseases taking immunosuppressive drugs, and in elderly and children (1–4). Two species are responsible for gastrointestinal infections: *Enterocytozoon bieneusi* and *Encephalitozoon intestinalis* (5). Both microsporidia are transmitted by the oral route and cause abdominal cramping, diarrhea, malabsorption, and weight loss in patients with AIDS (6). *E. intestinalis* also infects and develops inside intestinal macrophages, allowing the infection to spread from the intestine to other organs (7). A very limited number of drugs are available for treating intestinal microsporidiosis; these include albendazole and fumagillin, which are at least partially effective in reducing the parasite count (2,7). A strong immune response of an individual is mainly responsible for controlling the pathogen. The compartmentalized response against *E intestinalis* infection is primarily mediated by CD8^+^ and CD4^+^ T cells together with interferon (IFN) and interleukin (IL)-12 cytokines (8,9).

The intestinal mucosa (IM) acts as a host to a variable microflora, which plays an important role in nutrition absorption and immune function, among others. The immune response of the IM plays a critical role in maintaining the commensal homeostasis and protecting the host against pathogens (10,11). The immune response of IM against *E. cuniculi* infection is majorly mediated by antigen-specific intraepithelial lymphocytes (IELs) (12). Considering the entry point of microsporidia to be intestinal mucosa, there is a lack of knowledge of local immunity, especially in individuals on immunosuppressive medications.

The past decade has witnessed a surge in the use of immunosuppressive drugs for the treatment of neoplastic and autoimmune disease patients, as well as in patients undergoing transplantation. Cyclophosphamide (Cy) is one such immunosuppressive drug that has been recommended by the World Health Organization (WHO) and is on the list of drugs most frequently used by the Public and Private Health Systems owing to its efficacy, cost-effectiveness, and lesser side-effects than other drugs with similar actions, such as dexamethasone (13). Cy is primarily used against autoimmune and alloimmune diseases (14,15), in treating patients undergoing transplantation, such as bone marrow recipients (16) and in cancer treatment (17,18).

Cyclophosphamide is a cytotoxic alkylating agent that binds to DNA; its major effects on the body include cellular apoptosis and myelosuppression, and decreased lymphocyte, neutrophil, red blood cell, and platelet count. However, it also possesses immunomodulatory effects, which have not yet been clarified, such as (i) expansion of antigen-specific T cells, (ii) expansion of T cell-specific cytokines (IFNs, IL-7, and IL-15), (iii) decrease in regulatory T cells (Treg), and (iv) increased mobilization of dendritic cells from bone marrow, with activation of intracellular machinery for antigen processing and presentation (19). Thus, this anticancer agent is known for inducing immunogenic cancer cell death, subverting immunosuppressive T cells, and promoting Th1 and Th17 cells that control cancer outgrowth (20, 21).

We have shown previously Cy to suppress the immune system of mice; intraperitoneal infection of mice with *E. cuniculi* resulted in a disseminated, acute, and fatal encephalitozoonosis. These results were associated with the immunosuppressive effects of Cy (22). This study was designed to describe how the immunosuppressive effects of cyclophosphamide compromise the intestinal immune response against *E. intestinalis*, one of the most prevalent microsporidia in opportunistic infections in humans. Herein, we show a higher number of CD8^+^ and CD4^+^ T lymphocytes in IM in association with pro-inflammatory cytokines to be responsible for resolution of *E. intestinalis* infection in both Cy immunosuppressed and immunocompetent mice, despite the immunosuppressive activity of Cy observed in cells populations of Peyer’s plaques, mesenteric lymph nodes, peritoneum and even part of the immune population of the IM.

## Results

### Immunosuppressed mice showed transitory enteritis with fast resolution

All infected mice survived the oral infection by *E. intestinalis*, with no evidence of symptoms and macroscopic lesions in the gut. Microscopically, lymphoplasmacytic enteritis was observed in infected animals (*infected* and *Cy-infected)*, mostly affecting the duodenum and ileum, especially at 7 and 14 dpi (Fig. 1a, b). In addition to the inflammatory infiltrate, mucosal ulceration caused exposure of the underlying lamina propria (Fig. 1b), apical and mural multifocal necrosis, and villus-based epithelial proliferation. At 21 and 28 dpi, young cells and an increased number of mitoses were observed in the small intestine, suggesting a restoration of tissue integrity, via resolution of the inflammatory process and tissue remodeling. *E. intestinalis* spore clusters were observed on the glandular base (Fig.1c), with a higher fungal burden observed in the *Cy-infected* than the *infected* (Fig. 1d) group. However, in both groups, the fungal burden and the lesions progressively decreased from 7 to 28 dpi (Fig. 1d). No *E. intestinalis* spores were observed in uninfected mice. Thus, infection by *E. intestinalis* via oral route caused lymphoplasmacytic enteritis, accompanied by favorable proliferation in both immunosuppressed and non-immunosuppressed groups; however, the fungal burden was higher in the Cy-treated mice. Macroscopically, the infection led to the enlargement of the spleen and intestinal lymph nodes in the infected animals.

**Figure 1.**
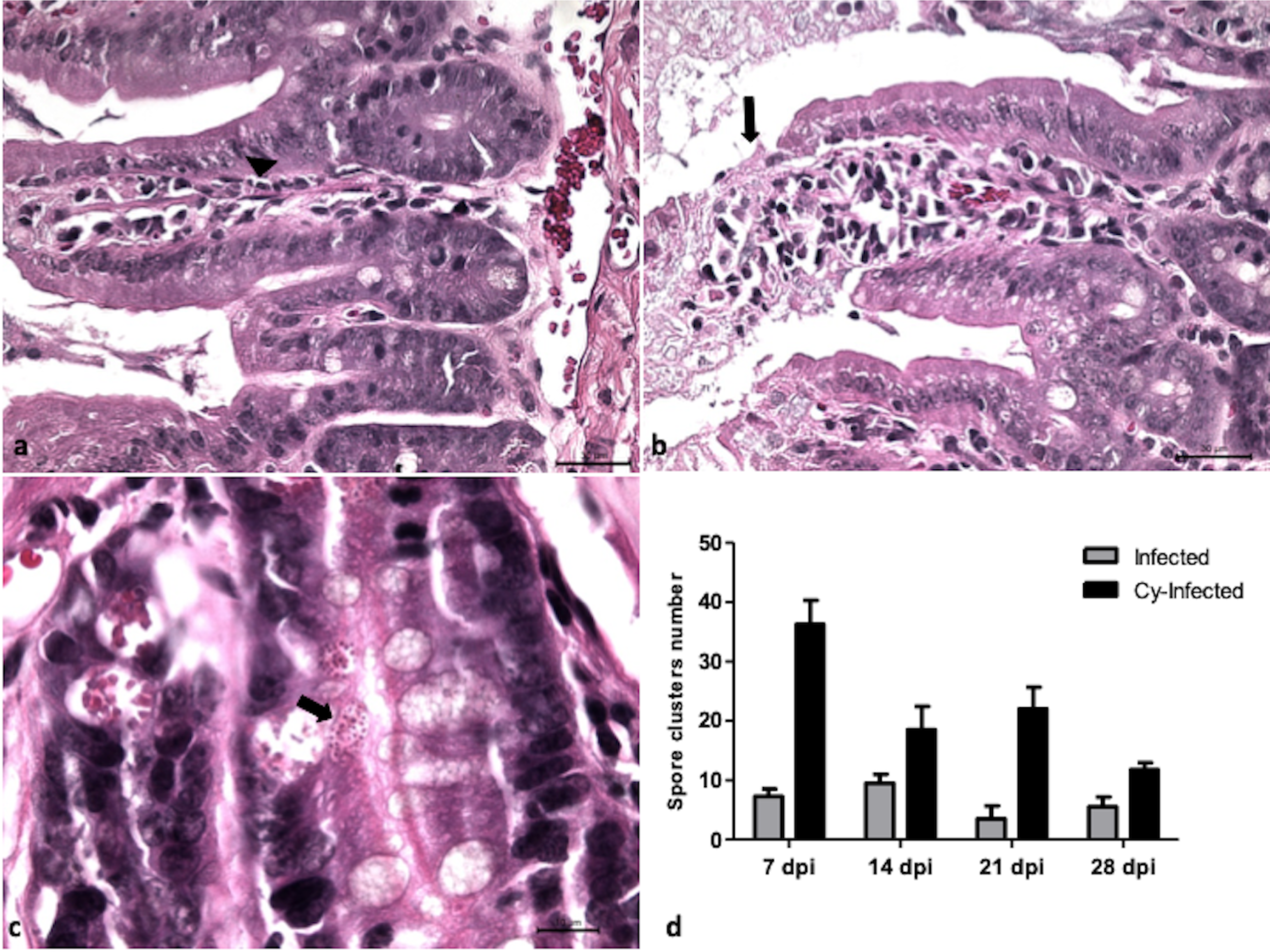
a) Photomicrograph of lymphoplasmacytic enteritis in the small intestine (arrowhead) in *Infected* mice. b) Ulceration (arrow) of intestinal mucosa of *Cy-Infected* mice. c) Clusters of *E. intestinalis* (arrow) at the glandular region in the mucosa of the ileum of *Cy-Infected* mice. d) Spores counting in the *Infected* and *Cy-Infected* mice. HE staining. ANOVA test with Tukey’s posttest showed p <0.01** and p <0.001***.

### CD8^+^ T lymphocytes increased in intestinal mucosa in infected mice

The *infected* and *Cy-infected* groups showed a significant increase in CD8^+^ T lymphocyte population as compared to controls at 7, 14, and 28 dpi (Fig. 2). This is in line with a previous study that demonstrated CD4^+^ T and CD8^+^ T lymphocyte subpopulations to play a substantive role in protecting against peroral infection of *E. intestinalis* (8). Another study reported a significant increase in the CD8αα subset of IELs in response to oral *E. cuniculi* infection (12).

**Figure 2.**
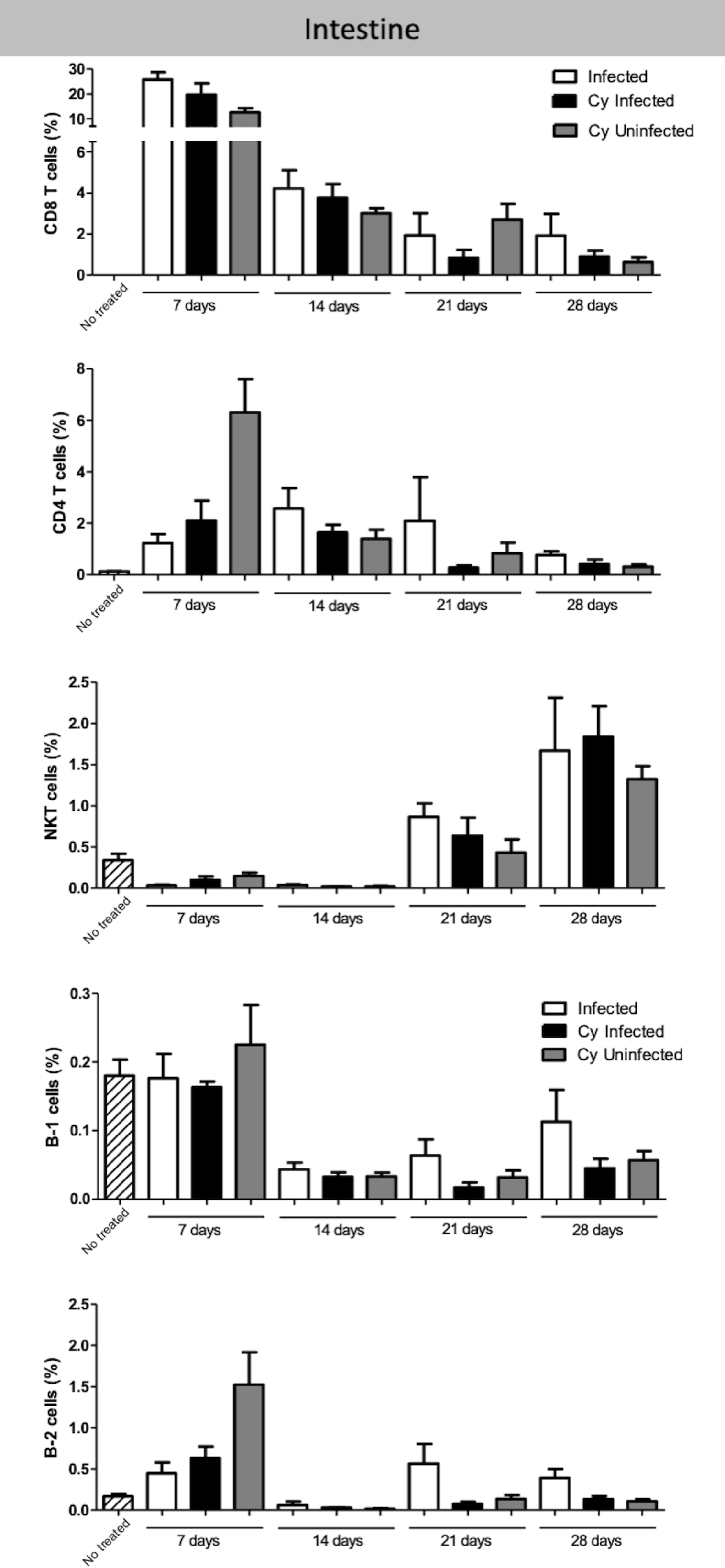
Evaluation of T and B cell population in the intestinal mucosa of the mice infected with *E. intestinalis* and treated or not with Cy at 7, 14, 21 and 28 dpi. Percentage of lymphocytes CD8 T (CD19^−^ CD4^−^CD8^+^), CD4 T (CD19^−^ CD8^−^ CD4^+^), NKT cells (CD19^−^ CD4^+^ NK1.1^+^), B-1 (CD23^−^ CD19^+^) and B-2 cells (CD23^+^ CD19^+^). ANOVA test with Tukey’s post-test showed p < 0.05*, p <0.01** and p <0.001*** compared to non-infected controls.

Immunosuppressive therapy with Cy could not cause a significant difference in CD8^+^ T lymphocyte population among the infected mice. Moreover, CD8^+^ T lymphocyte population witnessed a 10-time decrease, when compared 7 with 28 dpi (Supplementary Fig. 1). Together, the results showed that CD8^+^ T lymphocyte peak was observed at 7 dpi and was associated with a high number of *E. intestinalis* spores and lymphoplasmacytic enteritis. At the same time, there was a gradual reduction in the CD8^+^ T cell population, fungal burden, and histological lesions in infected (*infected* and *Cy-infected*) animals at 14, 21, and 28 dpi, indicating the resolution of infection in both immunosuppressed and non-immunosuppressed groups. The treatment with Cy resulted in a significant increase in the CD4^+^ T lymphocyte population in the uninfected group. In addition, *infected* and *Cy-infected* groups also showed an increase in this population at 14 and 28 dpi as compared to the untreated control (Fig. 2), suggesting that *E. intestinalis* stimulated CD4^+^ T expansion.

Further, the NKT cells decreased significantly in the *infected*, *Cy-infected*, and *Cy-uninfected* groups as compared to the *uninfected* control at 7 and 14 dpi (Fig. 2). However, these cells increased in the *infected* and *Cy-infected* groups in the later stages of infection (Supplementary 1). A significant reduction in B-1 cells was reported in the *infected*, *Cy-infected*, and *Cy-uninfected* groups as compared to the *uninfected* control at 14, 21, and 28 dpi (Fig. 2). Moreover, B-2 cells decreased in the *infected* and *Cy-infected* groups as compared to the *uninfected* control at 14 dpi but increased significantly in the *infected* group at 28 dpi (Fig. 2). Another notable observation was a decrease in the dendritic cells in PP in the *infected* and *Cy-infected* groups (Fig. 3a). PP in the *infected* group was more evident, with a marked increase in the germinal center activity, thereby expanding the lymphoid center (Fig. 3b). Moreover, PP from *Cy-infected* mice showed a rarified lymphoid tissue (Fig. 3b), enlarged lymphatics, and clusters of *E. intestinalis* spores (data not shown).

**Figure 3.**
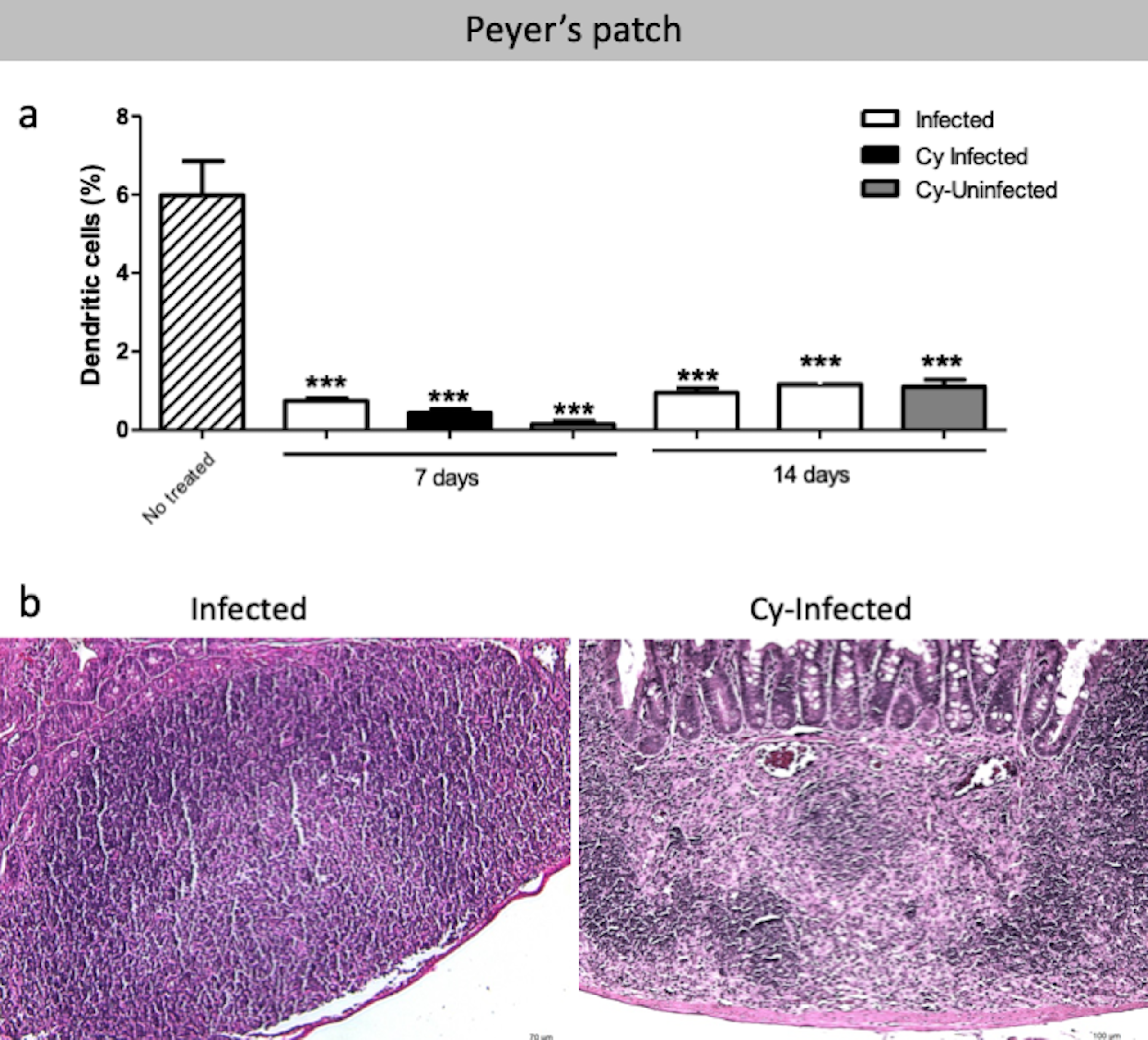
Peyer’s patch analyzed of the mice infected with *E. intestinalis* and treated or not with Cy for 7, 14, 21 and 28 dpi. a) Percentage of dendritic cells (CD11c^+^) in Peyer’s Patch. b) Photomicrograph of Peyer’s Patch with lymphoid expansion in *Infected* group or rarified lymphoid tissue in *Cy-Infected* group. HE staining. ANOVA test with Tukey’s post-test showed p <0.05*, <0.01** and <0.001*** compared to non-infected controls.

Interestingly, Cy treatment (*Cy-uninfected*) showed an immunomodulatory effect on lymphocyte populations in the intestinal mucosa. Overall, elevated levels of CD8^+^ T, CD4^+^ T, and B-2 cell populations were observed as compared to other groups (Fig. 2).

### T and B lymphocytes decreased in mesenteric lymph nodes of *Cy-infected* mice

The populations of B-2, CD4^+^ T, CD8^+^ T, and NKT cells decreased at 21 and 28 dpi in the *Cy-infected* group as compared to the *infected* group, suggesting the immunosuppressive effect of Cy. At 14 dpi, a higher percentage of macrophages was present in the *Cy-infected* group as compared to the *infected* group; however, this witnessed a decrease at 21 and 28 dpi (Fig. 4).

**Figure 4.**
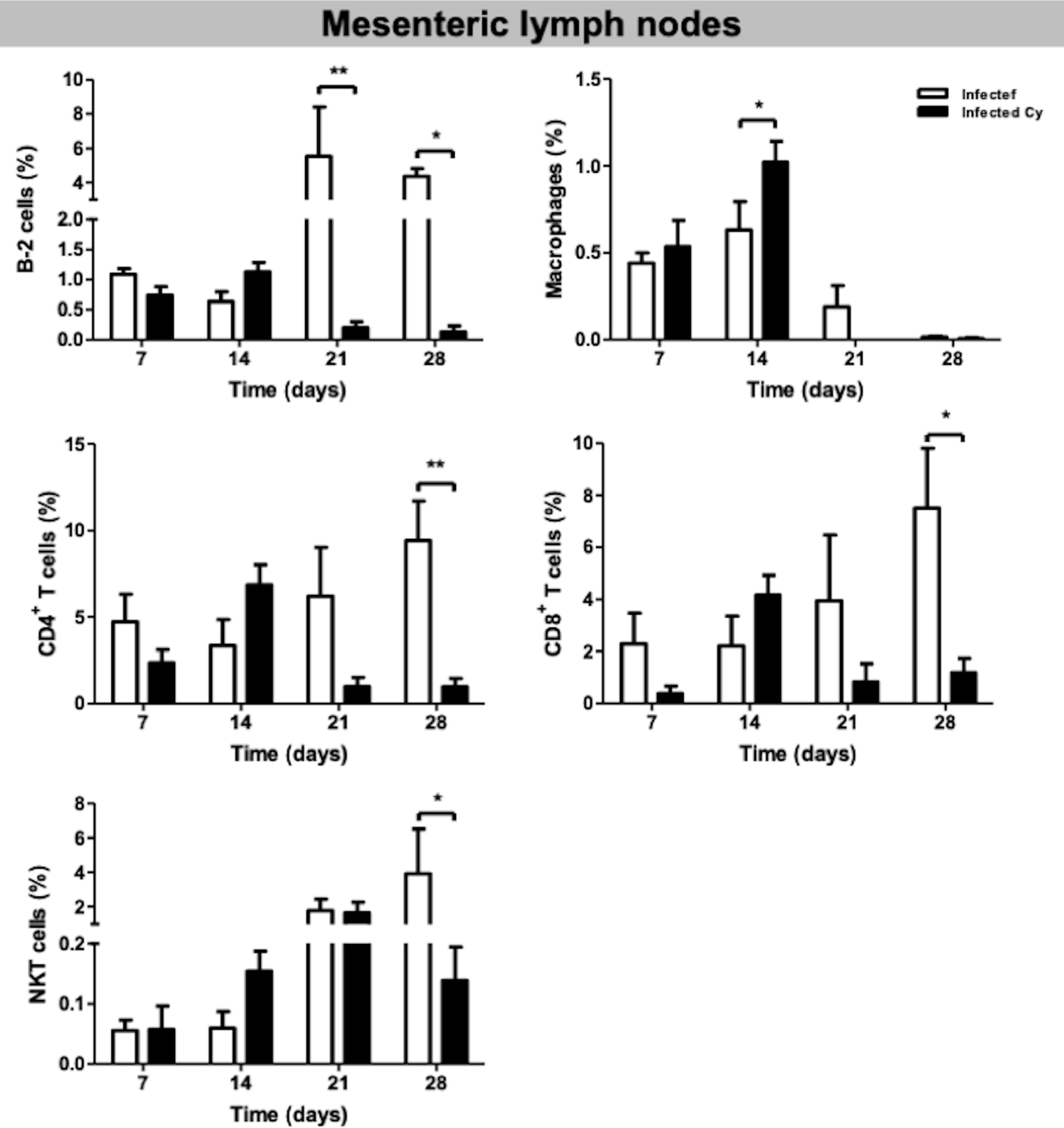
Cell population in the mesenteric lymph nodes of the mice infected with *E. intestinalis* and treated or not with Cy for 7, 14, 21 and 28 dpi. Percentage of B-2 cells (CD23^+^ CD19^+^), macrophages (CD19^−^ CD11b^+^), CD4 T cells (CD19^−^ CD8^−^ CD4^+^), CD8 T cells (CD19^−^ CD4^−^ CD8^+^) and NKT cells (CD19^−^ CD4^+^ NK1.1^+^). ANOVA test with Tukey’s post-test showed p <0.05* and p <0.01**.

### T and B lymphocytes decreased in the peritoneum of *E. intestinalis-*infected mice

The CD4^+^ T, CD8^+^ T, and NKT cells decreased significantly in all *infected* groups as compared to the *uninfected* control (Fig. 5). Moreover, CD8^+^ T, B-1, and B-2 lymphocytes decreased significantly in all Cy-treated groups at 7 and 28 dpi (Fig. 5). At 14 dpi, both B-1 and B-2 cell populations and macrophages increased in the *infected* group as compared to the *Cy-infected* group (Fig. 5). However, the macrophage number decreased in the *infected*, *Cy-infected*, and *Cy-uninfected* groups at 14 and 28 dpi (Fig. 5). Overall, the infected mice showed a decreased number of B-1, B-2, CD4^+^ T, and CD8^+^ T cells, with a statistical difference at 14 dpi (Supplementary Fig. 2). This difference was maintained at 21 and 28 dpi for B cells, but not for T cells.

**Figure 5.**
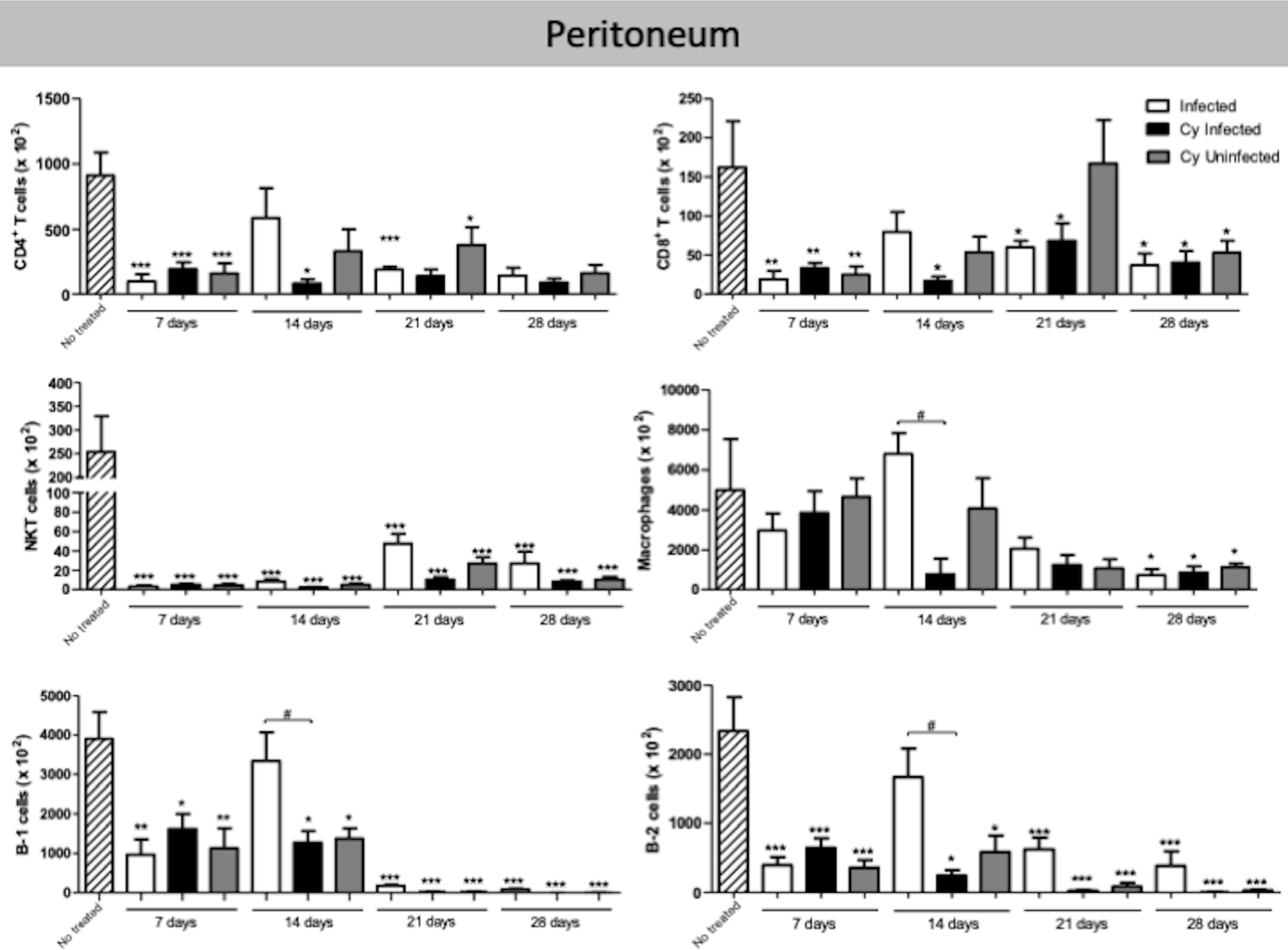
T and B population in the peritoneum of the mice infected with *E. intestinalis* and treated or not with Cy for 7, 14, 21 and 28 dpi. The number of CD8 T cells (CD19^−^ CD4^−^ CD8^+^), CD4 T cells (CD19^−^ CD8^−^ CD4^+^), NKT cells (CD19^−^ CD4^+^ NK1.1^+^), B-1 cells (CD23^−^ CD19^+^), B-2 (CD23^+^ CD19^+^) cells, and macrophages (CD19^−^ CD11b^+^). ANOVA test with Tukey’s post-test showed p <0.05*, p <0.01** and p <0.001*** compared to non-infected controls.

### Pro- and anti-inflammatory cytokines detected in the ileum of infected mice

*E. intestinalis* infection was associated with increased levels of various pro- and inflammatory cytokines, including IL-10, IL-17a, TNF-α, IFN-γ, IL-6, IL-4, and IL-2 at 14 dpi (Fig. 6). High levels of TNF-α persisted at 21 and 28 dpi, whereas other cytokines were not detected at these dpi (Fig. 6). These results implied that Cy treatment increased the production of IL-10 and IFN-γ at 7 dpi in the *Cy-uninfected* group as compared to the *uninfected* group. The highest level of cytokines was observed in the *Cy-infected* mice and included mostly IL-2, IL-6 and IL-4. The cytokine IL-17a was detected in all groups in all experimental animals although no statistical difference was observed between them. Moreover, the *uninfected* group showed enhanced levels of TNF-α (Fig. 6). Overall, the cytokines levels in the present study after *E. intestinalis* infection corroborate with the histopathological results obtained in all *infected* groups at 7 and 14 dpi, reflecting the immune system-mediated attempt to counter the intestinal infection.

**Figure 6.**
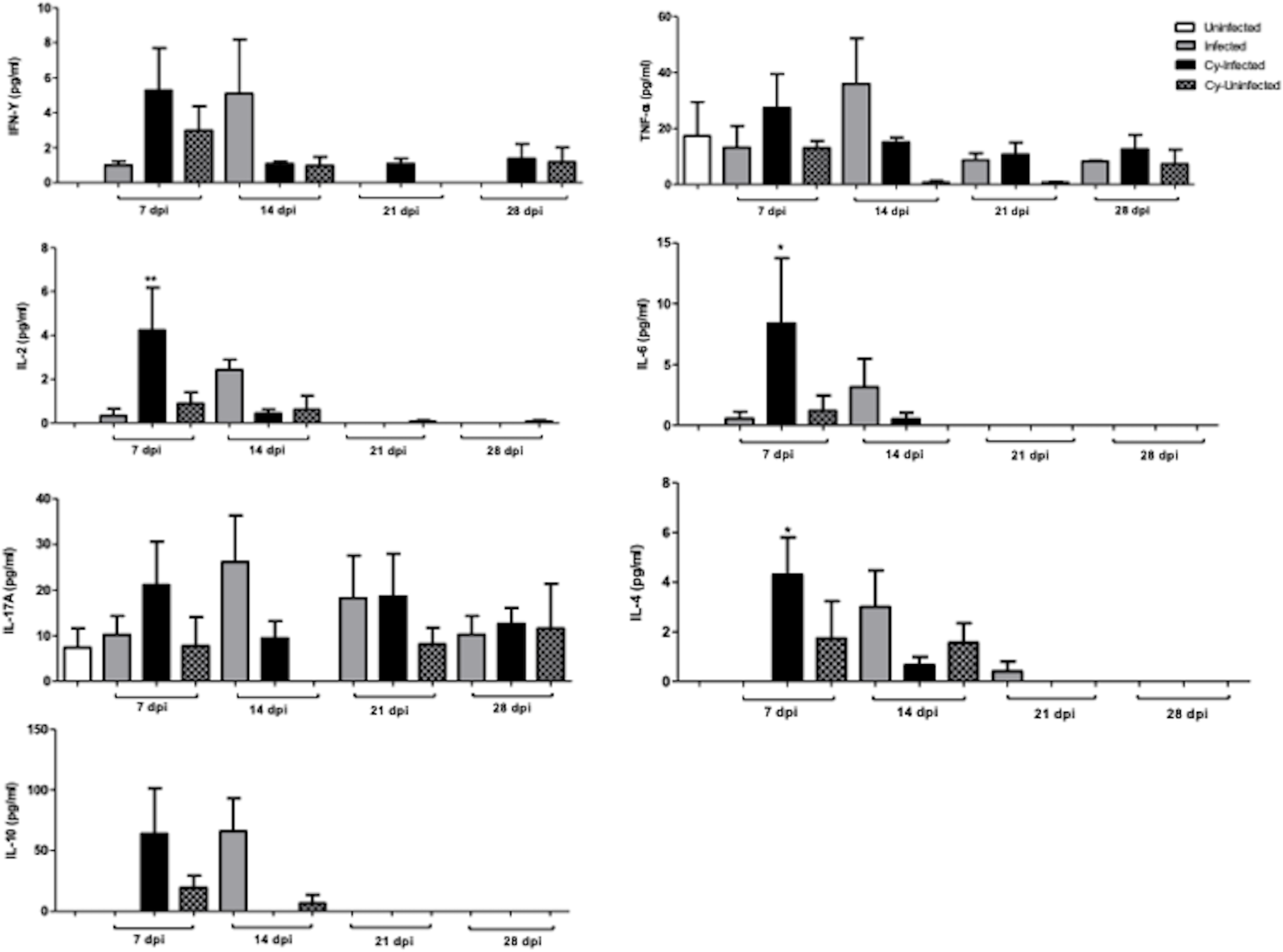
Proinflammatory cytokines (IL-2, IL-6, IFN-γ, and TNF-α) and anti-inflammatory cytokines (IL-4, IL-10, and IL-17A) levels in the intestine of mice infected with *E. intestinalis* and treated or not with Cy for 7, 14, 21, and 28 dpi. ANOVA test with Tukey’s posttest.

### Transmission electron microscopy

Transmission electron microscopy (TEM) results showed *E. intestinalis* spores of varying shapes, from oval to piriform, to be present close to the microvillosities and in the cytoplasm of enterocytes (Fig. 7) of the *Cy-infected* group. Additionally, two patterns of the morphological invasion were observed: endocytosis (Fig. 7 a,b) and injection of sporoplasm by polar tubule into the host cells (Fig. 7 c,d). The discontinuous microvillosities, present next to spores, showed invaginations, manifesting their attempt to engulf spores, a mechanism similar to phagocytosis. However, no phagocytic vacuoles were observed (Fig. 7 a,b,c). The cytoplasm surrounding the spores was found to be electrodense with a high number of mitochondria (Fig. 7 a,b,c). Moreover, surrounding the *E. intestinalis* sporoplasm injection demonstrated a loss of microvillosities and cytoplasmic membrane projections involving the extruded polar tubule that appeared to form a channel for its entry into the electrodense cytoplasm (Fig. 7 d).

**Figure 7.**
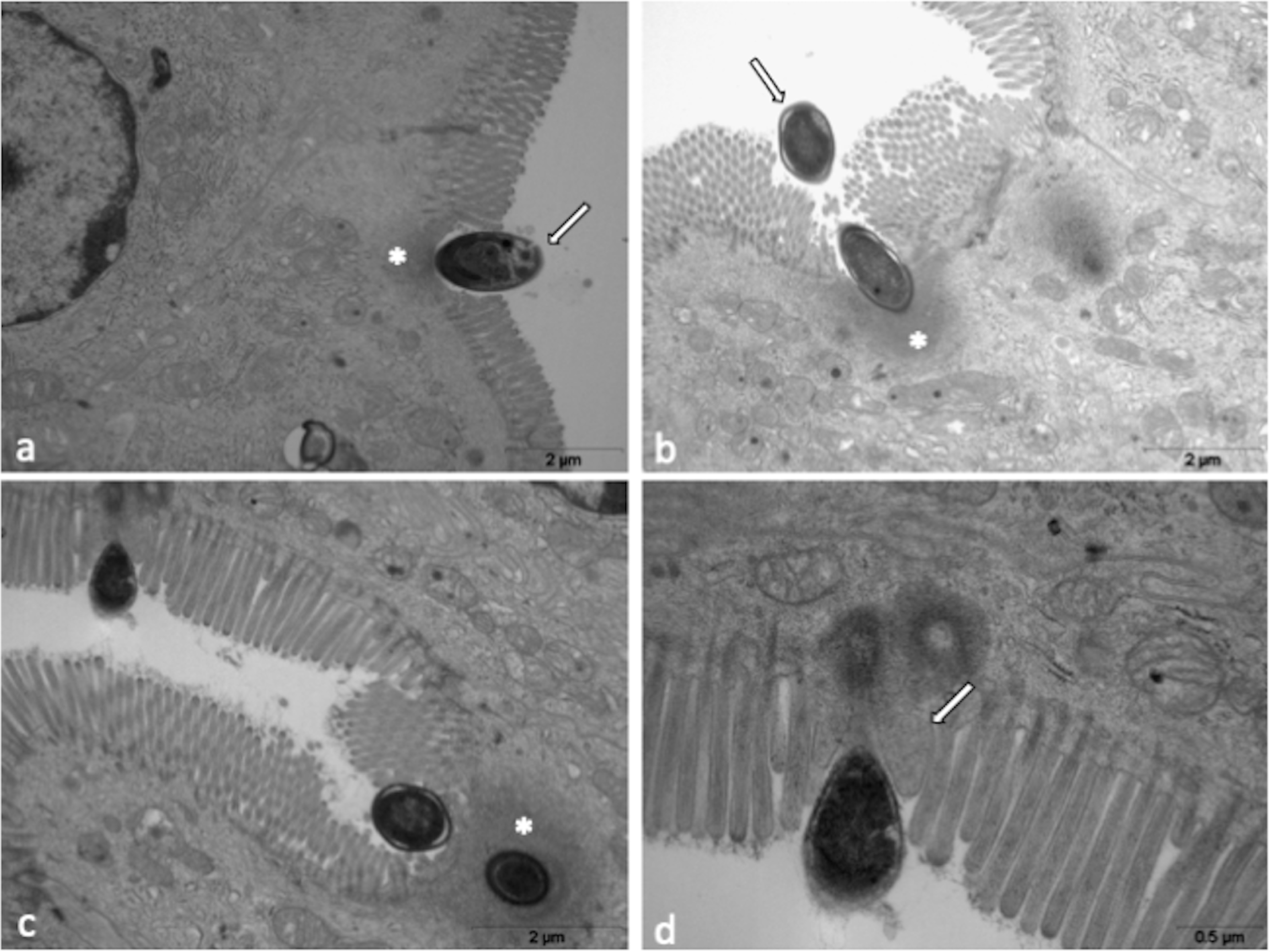
TEM from the intestine of *Cy-Infected* mice. a, b, and c*) E. intestinalis* spores adhered to and enterocyte (*). Note that microvillosities disappeared, the membrane shows invaginations associated with an electrodense area in the cytoplasm close to the spore (*). d) Injection of the sporoplasm with loss of microvillosities (arrow). Projections of cytoplasmic membrane involving the extruded polar tubule (arrow).

## Discussion

*E. intestinalis* invades and develops in the cytoplasm of enterocytes, causing persistent diarrhea in humans with AIDS (7) or in individuals immunocompromised by cancer or chemotherapy (23,24). Microsporidia also infect the macrophages of the lamina propria that help in its dissemination to kidneys and the hepatobiliary tract (25). In the present study, we observed oral infection with *E. intestinalis* not to be associated with clinical signs, such as diarrhea, even in the Cy-immunosuppressed mice. However, microscopically, lymphoplasmacytic enteritis was observed in association with *E. intestinalis* spores.

The adaptive immune response is very crucial for containing microsporidiosis. For example, T cells-deficient mice (athymic/nude or SCID) are incapable of successfully killing the pathogen and die due to encephalitozoonosis as opposed to immunocompetent mice (1,26). Similarly, IFN-γ^−/−^mice lacking major proinflammatory cytokines showed hepatomegaly, colicystitis, splenomegaly, intestinal enlargement, and ascites three to four weeks post infection with *E. intestinalis*. Moreover, these mice could live only up to six weeks post infection (25). Therefore, physical or genetic deletion of immune cells or cytokines represents a situation that does not correspond to clinical observations observed in individuals immunosuppressed by drugs. Since immunosuppressive drugs act on different cells of the immune system, leading to a diversity of immunocompromised states, we used Cy to mimic the physiological immunosuppression situations.

We observed that *Cy-infected* mice showed a higher number of spores, reflecting the increased susceptibility to pathogen upon treatment with Cy, as previously reported by our group in experimental *E. cuniculi* infection in mice (22). Unexpectedly, the present results showed a decrease in fungal burden and histopathological lesions in immunosuppressed mice at 21 and 28 dpi, indicating successful elimination of pathogen. Cy treatment is lymphotoxic and rapidly decreases the T and B cell populations (14). However, a study reported it to cause an extensive mobilization of immune cells from the bone marrow and other lymphoid organs (18). In the present study, we observed an increase in CD4^+^ T and CD8^+^ T cells population in the IM of Cy-treated mice. These results may be associated to two possible explanations: i) the displacement of immune cells from nearby immune sites such as Peyer’s patches, lymph nodes and peritoneum favored the assembly of an immune response in the gut capable of controlling infection, however, in this case, it should be considered that such reduction could be linked to the suppressive effect of Cy on these immune sites, or ii) low action on the immune population of the intestinal mucosa, especially T cells, data that may be linked to a possible immunomodulatory activity of Cy, which should be further explored

Another observation that point toward resolution of infection is the increased presence of mitosis in the intestinal crypts of the small intestine associated with enteritis in all infected animals, indicating tissue remodeling in the intestine under both pathological and physiological conditions (27). Since enterocytes serve as the first physical barrier against pathogens in intestine, maintenance of a strong epithelial barrier is crucial for normal physiology and evasion of infections. Moreover, epithelial cells secrete important chemoattractants that are involved in cell chemotaxis (28). The IELs constitute a sub-population of T lymphocytes in the IM that has been implicated in both intestinal homeostasis and inflammation, as observed in intestinal toxoplasmosis. Another important category of immune cells is dendritic cells that reside in the lamina propria beneath the IM and act as antigen presenting cells. They play a crucial role in containing toxoplasmosis. Lamina propria also comprises CD4^+^ T, CD8^+^ T, and B lymphocytes; B cells are known for producing secretory IgA that is responsible for the exclusion of environmental antigens (29).

There are reports stating an effective immune response against microsporidia to involve majorly CD8^+^ T cells (30,31). Interestingly, other studies report CD8^+^ T cells and not CD4^+^ T cells to be indispensable in successfully eliminating pathogens in intraperitoneal infection by *E. cuniculi* (30,32). In fact, the dichotomic role of CD4^+^ T cells has been shown to be associated with the infection route. It has been shown that the oral infection with *E. cuniculi* stimulates a synergistic effect of CD8^+^ and CD4^+^ T cells (12). The present study also reported an increased number of CD4^+^ T cells in the intestines of infected mice, contributing to successful *E. intestinalis* elimination. We believe this increase in CD4^+^ T cells to be attributed to the immunomodulatory effect of Cy in the intestinal mucosa.

Evidence from literature reports CD4^+^ and CD8^+^ T cells, associated with IFN-γ and IL-12 cytokines, to be important players in eliminating *E. intestinalis* infection (8,9). The present results reinforce this hypothesis; as evident by a significant increase in CD8^+^ T cells in all infected groups. A subsequent decrease in CD8^+^ T cell population was associated with a decrease in spore numbers and histopathological lesions. At 7 and 14 dpi, the increased production of TNF-α, IFN-γ, and IL-10 cytokines was noticed in the ileum of infected mice, together with an increase in IL-2 and IL-4 in *Cy-infected* mice. The results suggest that besides IFN-γ (8,9), other cytokines, including anti-inflammatory cytokines, play a crucial role in mediating an intestinal immune response against *E. intestinalis*, for the resolution of enteritis and pathogen killing.

The IELs comprise a heterogeneous population, predominantly composed of CD8^+^ T (CD8αα and CD8αβ) cells and a few CD4^+^ T lymphocytes and known to play an important role in oral infections. Upon infection, an expansion in the number of these cells causes increased production of IFN-γ, cytolytic properties of which inhibit intestinal *E. cuniculi* proliferation and dissemination (12). In fact, our results showed an increased CD8^+^ T IELs in intestinal *E. intestinalis* infection. However, CD8^+^ T cells decreased in the peritoneal cavity, suggesting their migration to the site of pathogen proliferation. This finding is in corroboration to previous studies that have already shown cells in the peritoneal cavity to migrate to other sites (33,34).

Our group has previously shown B-1 cells to be an important player in the generation of the immune response against *E. cuniculi* infection (35,36). B-1 cells are important in generating adaptive immune response and antibody production, which is independent of T cells (37). However, B-1 cells are also dependent on T cells for generating an effective immune response, such as cell-mediated hypersensitivity (38,39) and rejection of aloenxerts (40). Moreover, adoptive transfer of B-1 cells activates T cells that produce IFN-γ (41). Here, we showed a decrease in B-1 cell frequency in *infected* animals as compared to controls of both intestine and peritoneal cavity. We hypothesized that B-1 cells from the peritoneal cavity differentiated into B-1 cell-derived phagocyte (B-1 CDP) in infected mice, which, in turn, promoted phagocytosis of *E. intestinalis* spores. *In vitro* studies showed that B-1 cells may also differentiate into mononuclear phagocytes, which upon attachment to a substrate, acquire a myeloid phenotype (33). Moreover, *Propionibacterium acnes* infection of B-1 cells of myeloid lineage induced only differentiation into phagocytes (42); however, the role of B-1 cells in the intestines warrants further understanding of microsporidia infection.

We also found a decrease in the NKT cells in the intestine and mesenteric lymph nodes in all infected mice. NKT cells are known to play a protective role in *Toxoplasma gondii* oral infection although these cells are susceptible to direct invasion by the parasite (43). In fact, NKT cells develop a hypermotility phenotype *in vivo* during *T. gondii* oral infection, suggesting manipulation of motility of immune cells by *T. gondii*, which assist in the spread of the causative organism (44,45), as already shown in macrophages (45). *E. cuniculi* uses macrophages as a Trojan horse (46); however, whether this phenomenon occurs with NKT cells in microsporidiosis remains largely unknown.

While an effective immune response against microsporidia is predominantly driven by T cells, dendritic cells have also been shown to play an important and critical role in stimulating these cells. Dendritic cells of the IM act as antigen-presenting cells and are responsible for priming T naive cells in effectors and memory cells upon infection (47). The secretion of cytokines and chemokines from infected enterocytes results in migration of these cells from PP to the mucosa (28). On the other hand, a study reported *in vitro* inhibition of dendritic cell differentiation by *E. intestinalis* (48). On the same lines, we, in the present study, demonstrate a reduction in dendritic cells in PP of infected animals at 7 and 14 dpi, which was associated with a higher number of fungal spores and histopathological lesions, suggesting the migration of these cells to the site of infection. In the gastrointestinal tract, dendritic cells play an important role in suppression of colitis development, inducing the traffic of T regulatory cells in the intestine, and inducing IgA secretion from B cells in the intestine (28). These are also involved in inducing oral tolerance and preventing inflammatory response mediated by gut microbiota and ingested antigens (28). The breach in this function may decrease the number of T regulatory cells and increase the number of cells producing *Th1* and *Th17* cytokines (28). A lower frequency of dendritic cells observed in the present study is related to a higher parasite burden; however, this is yet to be clarified.

It has been previously shown that microsporidia spores infect new cells mostly by injection of sporoplasm into the host cells via the polar tubule (49). However, different cell lineages are capable of phagocytosing microsporidia spores (50,51), suggesting endocytosis to be an important mechanism in intestinal microsporidiosis. In order to evade this protective mechanism, microorganisms have developed complex systems to subvert endocytosis by host cells, allowing invasion of even those cells that generally do not phagocytose (52). Previous studies have suggested microsporidia to induce invaginations of the cell membrane of host cells close to the polar tubule in a process similar to endocytosis, causing injection of sporoplasm into the host cell. The results of TEM showed that a phenomenon similar to phagocytosis occur more frequently in *E. intestinalis* infection.

The results of this study showed that the increase in the CD8^+^and CD4^+^ T cell population in the intestinal mucosa of Cy immunosuppressed mice and orally infected with *E. intestinalis*, in association with the presence of pro-inflammatory cytokines, controlled the infection by the opportunistic fungus, although reduction of cellular populations at immune sites has been observed, confirming a state of immunosuppression, reinforcing that the selective effect of Cy should be better understood.

## Methods

### Animals

Specific pathogen-free, 6–8-week-old C57BL/6 mice were purchased from the Federal University of São Paulo (CEDEME, UNIFESP), Brazil. Animals were housed under sterile conditions at the Animal Facility of Paulista University, São Paulo, Brazil, and given food and water *ad libitum*. All animal procedures were performed in strict accordance with the Paulista University Ethics Committee (project license no. 313/15).

### *Encephalitozoon intestinalis* cultivation and experimental infection

Spores of *E. intestinalis* were purchased from Waterborne Inc., New Orleans, LA, United States. These were cultivated in rabbit kidney cells (RK–13) in Dulbecco’s Modified Eagle’s medium (DMEM) supplemented with 10% fetal bovine serum (FBS), pyruvate, non-essential amino acids, and gentamicin followed by incubation at 37 °C and 5% CO_2_. Spores were collected from the supernatant, washed thrice in phosphate-buffered saline (PBS), and counted using a Neubauer chamber.

### Study design

Mice were divided into four experimental groups: *infected*, mice infected with *E. intestinalis*; *uninfected*, non-infected and non-treated mice; *Cy-infected*, mice treated with cyclophosphamide and infected with *E. intestinalis*; and *Cy-uninfected*, mice treated with cyclophosphamide. The Cy-treatment protocol was previously established (22) and consisted of intraperitoneal injection of 100 mg/kg twice a week (Genuxal; Asta Medica Oncologia, São Paulo, Brazil). The treatment started at the day of infection until 28 days post infection (dpi). Mice were orally infected by gavage with 5× 10^7^ *E. intestinalis* spores. Non-infected mice served as control.

### Necropsy and tissue sampling

At 7, 14, 21, and 28 dpi, five animals from each group were euthanized with a mixture of ketamine (100 mg/mL), xylazine (20 mg/mL), and fentanyl (0.05 mg/mL). The samples of the intestine (duodenum, ileum), liver, kidneys, and lungs were collected and fixed in 10% buffered formalin for 72 h, routinely processed for histopathology, and stained with hematoxylin-eosin (HE) and Giemsa.

### Ultrastructural analysis by transmission electronic microscopy

Ileum samples of 1 mm thickness from Cy-infected mice were fixed in 2% glutaraldehyde in 0.2 M cacodylate buffer (pH 7.2) at 4 °C for 10 h. These were then fixed in buffered 1% OsO4 for 3 h. Subsequently, the samples were embedded in EPON resin, sliced into semi-thin cuts, and stained with toluidine blue. Then, ultrathin sections were double stained using uranyl and lead citrate and observed under the TEM LEO EM 906 at 80 kV at the Butantan Institute.

### Fungal burden

The paraffinized ileum was cut into 5-μm thick sections and evaluated histopathologically using HE staining to determine the fungal number. Fungal spores were counted randomly in at least 10 fields under the light microscope (40× objective magnification). The average number of spores from each mice was recorded and statistically analyzed.

### Phenotypic analysis of immune intestinal mucosal cells

Cells of the IM were obtained as previously described,^53^ with minor modifications. The small intestine was washed with 50 mL of Hanks’ balanced salt solution (HBSS)–2% FBS solution, longitudinally cut and separated into 2-cm segments. The mucosa was grated and submerged in HBSS–2% FBS solution supplemented with 0.1 M ethylenediaminetetraacetic acid (EDTA) at 37°C for 20 min. Then, samples were vortexed for 15 s and filtered using a cell strainer to remove the cell debris. Percoll gradient (70% and 40%) centrifugation was used to isolate the IM cells. After centrifugation, cells were washed with HBSS–2% FBS and resuspended in 100 μL of PBS–1% bovine serum albumin (BSA).

Mesenteric lymph nodes (MLN) and Peyer’s patches (PP) were isolated from the intestines using a scalped blade and washed in a cell strainer with HBSS–2% FBS. In addition, cells from the peritoneal cavity (PerC) were obtained by successive washes with at least 10 mL of HBSS–2% FBS. Finally, cell suspensions from PerC, MLN, and IM were washed with HBSS–2% FBS and resuspended in 100 μL of PBS–1%BSA. After centrifugation at 500 ×g for 5 min, each sample was incubated at 4 °C for 20 min with the anti-CD16/CD32 antibody. After incubation, cells were divided into two aliquots and resuspended in PBS–1%BSA, followed by incubation with monoclonal antibodies: APC-conjugated anti-mouse CD19, FITC, or PE-conjugated anti-mouse CD23, PerCP-conjugated anti-mouse CD4, FITC-conjugated anti-mouse CD8, APC-Cy7-conjugated anti-mouse CD11b, and PE-conjugated anti-mice CD11c (BD Pharmingen; San Diego, CA, United States). Finally, cell suspensions were run on the flow cytometer FACS Canto II (BD Biosciences; Mountain View, CA, United States). The cells were characterized according to their phenotypes in CD8 T (CD19^−^ CD4^−^CD8^+^), CD4 T (CD19^−^ CD8^−^ CD4^+^), NKT cells (CD19^−^ CD4^+^ NK1.1^+^), B-1 (CD23^−^ CD19^+^), B-2 cells (CD23^+^ CD19^+^) and dendritic cells (CD11c^+^); and analyzed with the software FlowJo (FlowJo LLC; data analysis software, Ashland, OR, United States).

### Cytokine quantification

For quantification of intestinal cytokines, 100 mg of ileum was sampled and treated with 1 mL of protease inhibitor (Sigma-Aldrich; St. Louis, MO, United States) at −80 °C. The sample was thawed and processed using a Precellys homogenizer for three cycles, 20 s each, filtering the homogenate using a cell strainer to remove the debris. Cytokines (IL-2, IL-4, IL-6, IL-10, IL-17, IFN-γ, and TNF-α) in the homogenate were measured using the CBA Mouse Th1/Th2/Th17 cytokine kit (BD Biosciences; CA, United States) according to manufacturer’s instructions. The kit consists of fluorescent beads coated with antibodies specific to cytokines. Briefly, 25 μL of each sample was added to capture beads and PE-labeled secondary antibodies. The samples were incubated for 2 h at room temperature in the dark, following which two-color flow cytometry analysis was performed on the FACS Canto II flow cytometer (BD Biosciences; Mountain View, CA, United States) and analyzed using the FCAP Array 1.0 software (BD Biosciences; CA, United States).

### Statistical analysis

Analysis of variance (ANOVA) tests and Tukey’s or Bonferroni’s multiple comparison post-tests were used for the statistical analysis of data. All values are expressed as mean ±standard error mean (SEM) with significance of *α* = 0.05 (*p* < 0.05). All graphs were generated using the “GraphPad Prism” version 5.0 for Windows (GraphPad Software Inc; La Jolla, CA, United States).

## Acknowledgments

We thank Magna Maltauro Soares (Instituto Butantan) for histopathological processing. We thank Fundação de Amparo à Ciência do Estado de São Paulo–Fapesp and Capes for financial support.

